# Rapid interannual reorganisation of Whiskered Tern *Chlidonias hybrida* breeding colonies within an urban wetland network

**DOI:** 10.64898/2026.10.09.757943

**Authors:** Andrei Daniel Cotoară, Mateusz Ledwoń, Relu Constantin Giucă, Emil Todorov, Alexandru-Cătălin Birău, Gergely Osváth

## Abstract

Colonial waterbirds breeding in unstable wetland habitats are expected to show flexible colony-site use, but how breeding activity redistributes within small networks of alternative nesting sites, particularly in urban wetlands, is poorly documented. We quantified the spatial organisation of breeding Whiskered Terns *Chlidonias hybrida* across nine lakes of Văcărești Natural Park, a spontaneously developed urban wetland in Bucharest, Romania, using nest counts repeated every one to three weeks over four breeding seasons (2022–2025). The number of occupied lakes fell from nine in 2022 to three in 2024, before a partial recovery to six in 2025, while the annual peak of simultaneously active nests ranged from 53 (2024) to 222 (2025). Spatial concentration at the annual peak survey was higher in 2024–2025 (dominant-lake proportion 0.55–0.56; Simpson’s concentration index C = 0.42–0.50) than in 2022–2023 (0.41–0.46; C = 0.29– 0.32), and the peak-survey distributions of 2024 and 2025 showed no overlap (Bray– Curtis dissimilarity = 1.0). The breeding season was long and asynchronous in 2022, when the largest colony shifted repeatedly among lakes, but compressed and synchronised in 2024–2025, when a single lake remained dominant throughout the season. Lake surface area (0.18–2.25 ha) was not detectably associated with occupancy or seasonal maximum nest numbers. These results indicate highly flexible colony-site use and rapid interannual reorganisation of breeding activity within a compact network of alternative nesting sites. Conservation and monitoring of urban wetland colonies should therefore encompass the entire network of potential nesting sites rather than only the currently dominant colony site.

## 1. Introduction

More than half of the world’s natural wetlands have been lost over recent centuries, with Europe among the regions most affected (Davidson, 2014). As natural wetlands disappear, habitats of anthropogenic origin, including those that emerge spontaneously within cities, play an increasingly recognised role in sustaining wetland biodiversity (Kowarik, 2011; Alikhani et al., 2021). Wetlands embedded in urban landscapes can support a disproportionately rich avifauna and provide breeding habitat for waterbirds (McKinney et al., 2011), yet colonial waterbirds, whose reproduction is concentrated at a small number of sites, remain comparatively little studied in urban settings.

For colonial waterbirds, the choice of a colony site is a key decision with major fitness consequences (Danchin et al., 1998; Barati et al., 2011). Where nesting habitat is stable, strong colony-site fidelity generally pays; where habitat is ephemeral or unpredictable, theory predicts weak site tenacity and flexible colony-site choice (McNicholl, 1975; Switzer, 1993). Empirical studies confirm that colonial larids and other waterbirds readily abandon sites after breeding failure, predation or disturbance (Burger, 1982; Nager et al., 1996), and that networks of discrete colony sites function as metapopulation-like systems with regular interannual exchange of breeders (Devlin et al., 2008; Schippers et al., 2009; Wilkinson and Jodice, 2022). Conspecific attraction and habitat copying, in contrast, tend to concentrate breeders at currently successful sites (Danchin and Wagner, 1997; Danchin et al., 1998), and site-occupancy dynamics track local environmental conditions (Wyman et al., 2014; Carrasco et al., 2017). The distribution of breeding activity among available sites therefore reflects a balance between dispersion and aggregation, whose benefits, such as predator swamping through synchronised breeding, are themselves environment-dependent (Ims, 1990; Descamps, 2019).

The Whiskered Tern *Chlidonias hybrida* is a colonial marsh tern that breeds on shallow, well-vegetated eutrophic water bodies, building floating nests anchored to floating-leaved or emergent vegetation (Cramp, 1985; Gochfeld et al., 2020). The extent and quality of floating macrophyte beds, governed largely by hydrological conditions, determine the availability of nesting habitat (Paillisson et al., 2006; Cieślińska et al., 2025), and abrupt water-level fluctuations are a major cause of nest loss and colony abandonment (Ortiz Lledó et al., 2018). Consistent with the instability of such habitats, marsh terns show weak colony-site fidelity: in the congeneric Black Tern *Chlidonias niger*, breeding-site abandonment is predicted by changes in vegetation and in local abundance (Wyman and Cuthbert, 2017). The Whiskered Tern has expanded rapidly in central and eastern Europe in recent decades (Ledwoń et al., 2014; Dayton et al., 2017; Keller et al., 2020), and Romania holds one of the most important European populations, with its stronghold in the Danube Delta, where colony dynamics are strongly driven by hydrological variability (Ion et al., 2026).

Most knowledge regarding colony-site dynamics in marsh terns derives from extensive natural or semi-natural wetland complexes (Ledwoń et al., 2014; Ion et al., 2026). Considerably less is known about how colonial species structure their breeding settlement within small, spatially heterogeneous urban wetlands, where the entire network of potential colony sites is compressed into a few tens of hectares. Văcărești Natural Park in Bucharest, Romania, provides a suitable system in which to examine these dynamics: an unfinished hydrotechnical reservoir abandoned in the early 1990s, spontaneously recolonised by wetland vegetation and wildlife, and designated as Romania’s first urban natural park in 2016 (Anastasiu et al., 2017). Although this mosaic of small waterbodies supports a rich urban breeding avifauna (Drăgan et al., 2024), including a regular breeding population of Whiskered Terns, the spatial organisation of its breeding waterbirds has not previously been studied.

Here we use four years (2022–2025) of systematic nest counts on nine lakes of Văcărești Natural Park to quantify the spatial organisation of breeding Whiskered Terns within this urban wetland. Specifically, we ask (1) how stable the distribution of breeding activity among lakes is between years, in terms of lake occupancy, the identity of the dominant colony lake and the overall concentration of nests; (2) how the spatial concentration of breeding changes within the season, and how closely the peaks of different colonies coincide in time; and (3) whether lake surface area is associated with the among-lake distribution of nests. Given the dependence of nesting habitat on unstable floating vegetation, we expected weak colony-site persistence and substantial interannual redistribution, combined with pronounced aggregation of breeders at a small number of sites at any one time. Throughout the paper, spatial organisation and reorganisation refer to the distribution of active nests among lakes; because individual birds were not marked, the movements of individuals are not addressed.

## 2. Materials and methods

### 2.1. Study area

The study was conducted in Văcărești Natural Park, an urban wetland complex located within the administrative boundaries of Bucharest, Romania. The park covers 183 ha on the site of a hydrotechnical reservoir abandoned in the early 1990s (Government of Romania, 2016), and its wetland habitats include extensive reedbeds, with at least 81 ha of *Phragmites*-dominated communities (APNV, 2023). The lakes are fed exclusively by precipitation and groundwater, with no channel connections or other external water supply, and water depth varies considerably among lakes (approximately 40–150 cm). Public access is restricted to designated trails, and fishing, hunting and bathing are prohibited within the park (APNV, 2023), so direct human disturbance at the lakes is low. The park comprises a mosaic of aquatic and terrestrial habitats, including several permanent and temporary water bodies that provide breeding habitat for a diverse waterbird community. The study focused on nine lakes within the wetland complex (Fig. 1) that were systematically surveyed for breeding Whiskered Terns between 2022 and 2025.

**Figure 1.**
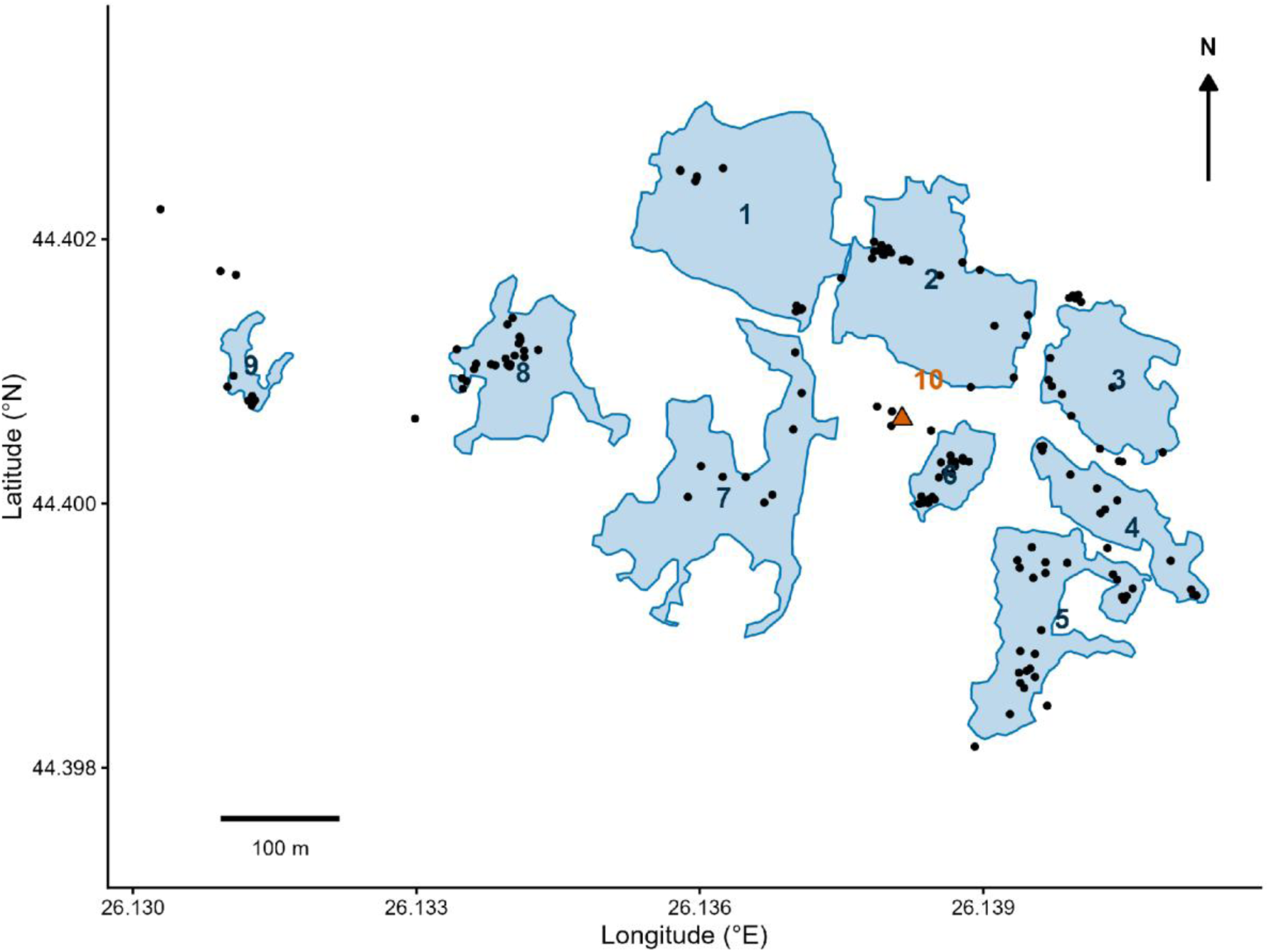
The nine monitored lakes in Văcărești Natural Park, Bucharest, Romania. Lake boundaries (blue polygons, numbered) were digitised from satellite imagery in Google Earth Pro; black points show all georeferenced active-nest observations (incubating adults, eggs or chicks) of Whiskered Terns *Chlidonias hybrida* from repeated surveys, 2022–2025. The orange triangle marks Lake 10, an additional small water body that could be monitored comparably only in 2025 (see Section 2.3).

All nine lakes were treated as distinct spatial units and monitored across all four study years, allowing interannual and seasonal comparisons of their relative use by breeding Whiskered Terns. Water levels fluctuate synchronously within two interconnected groups (Lakes 1–2 and 7, and Lakes 4–5), but independently across the isolated Lakes 3, 6, 8 and 9. The nine monitored lakes comprise the large majority of the open-water nesting habitat available in the park. Additional small pools along the survey routes were checked regularly and never held breeding Whiskered Terns, and most become completely overgrown by reed during the breeding season; one small reed-locked pond could not be inspected visually from the ground and remained unmonitored. Observations of the species were georeferenced wherever they occurred in the park, not only on the monitored lakes. Across the four seasons, only one record with confirmed breeding evidence (a single nest with eggs in July 2022, on a small pool northwest of Lake 9) and three records of possible breeding activity (nest building or territorial behaviour in 2025) were obtained away from the monitored lakes and Lake 10, together representing less than 0.1% of all nest-coded counts. The monitored network therefore held the overwhelming majority of the park’s breeding population.

### 2.2. Nesting substrates

Systematic macrophyte surveys were not conducted during the study, but opportunistic ground and drone photographs revealed high interannual variation in the nesting substrates used by Whiskered Terns (Fig. S3). In 2022, nests were located on floating mats of green algae *Cladophora* sp., debris islands of broken Common Reed *Phragmites australis* and, to a lesser extent, White Water-lily *Nymphaea alba*. Green algae remained a primary substrate in 2023. In 2024, nesting shifted to Water Crowfoot (*Ranunculus* sp., most likely *R. trichophyllus*) and *Rumex* sp. By 2025, nests on the dominant site (Lake 4) were built almost exclusively on amphibious Great Yellow-cress *Rorippa amphibia*. This sudden dominance reflects the species’ life history: its subterranean rhizomes can remain dormant during drawdowns, accumulating energy and expanding vegetatively when exposed to air (Akman et al., 2012). The extensive development of Great Yellow-cress on Lake 4 in 2025 coincided with the establishment there of the largest breeding aggregation of the study and may have provided particularly suitable nesting substrate. Nests were constructed from plant material commonly available in the immediate vicinity, including Rigid Hornwort *Ceratophyllum demersum*, Common Reed, Water Crowfoot and Great Yellow-cress. These observations were opportunistic rather than systematic and are therefore treated as ecological context, not as a formal environmental analysis.

### 2.3. Field monitoring

Breeding Whiskered Terns were monitored across four consecutive breeding seasons (2022–2025), with 10–19 surveys per season conducted at intervals of one to three weeks (Table S1). Surveys followed standardised transects and elevated vantage points to ensure complete visual coverage of all nine lakes. Active nests were counted per lake within a single day or, occasionally, over two consecutive days when required for full coverage (4 of 51 survey periods: three in 2022 and one in 2025; see Section 2.4), in which case the visits were merged and treated as effectively simultaneous. A nest was classified as active when an incubating adult, eggs or chicks were observed; nest building alone was not treated as evidence of an active nest. Surveys were conducted with binoculars (10 × 42) and a spotting telescope (20–60×). Counts represent the number of visually detectable active nests; detection probability was not estimated separately.

For each survey date, the number of active nests recorded on each lake was retained as a separate observation. Consequently, the dataset consisted of repeated observations of the same nine lakes throughout each breeding season.

An additional small water body (Lake 10) supported breeding in 2022 (up to three active nests) and 2025 (up to 28 active nests), but it could not be monitored consistently across years and was therefore excluded from all interannual comparisons.

### 2.4. Data processing

Total breeding abundance for each survey period was calculated as the sum of active nests across all nine lakes. Annual peak abundance was defined as the highest survey-level total recorded in a given season, forming the basis for interannual comparisons of spatial concentration and dissimilarity. To ensure spatial coverage without double-counting, complementary visits on consecutive days were merged into a single survey period using the maximum count per lake; these counts were treated as contemporaneous. Lake occupancy was defined as ≥ 1 active nest during a survey period, with the lake supporting the most nests identified as the dominant lake.

To quantify the spatial concentration of breeding activity, we calculated the proportion of all active nests occurring on the dominant lake during each survey period as:

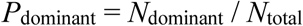

where *N*_dominant_ is the number of active nests on the lake supporting the largest aggregation and *N*_total_ is the total number of active nests recorded across all nine lakes during the same survey period.

To characterise nest distribution among lakes beyond the dominant lake, we quantified spatial concentration and evenness for each survey period with ≥ 10 active nests (to avoid small-sample instability) and for the simultaneous annual peak survey; equivalent metrics based on lake-specific seasonal maxima are provided in the Supplementary Material (Tables S2a, S2b and S3). Let *p*_i_ denote the proportion of active nests on lake *i*. Spatial concentration was quantified using Simpson’s concentration index (C):

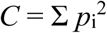

(Simpson, 1949); its reciprocal, 1/C, expresses the effective number of lakes used. Evenness among occupied lakes was quantified using Pielou’s evenness:

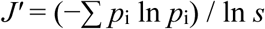

where *s* is the number of occupied lakes (*s* > 1; Shannon, 1948; Pielou, 1966). *J’* was not calculated when *s* = 1. Higher values of *J’* indicate an even distribution, whereas higher C values indicate stronger concentration. Because *s* varied among years, *J’* was interpreted alongside C. Index terminology follows Magurran (2004).

Interannual shifts in spatial distribution were quantified using Bray–Curtis dissimilarity (Bray and Curtis, 1957) between relative lake distributions at annual peak surveys:

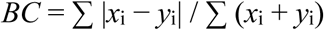

where *x*_i_ and *y*_i_ are nest proportions on lake *i* in two compared years. Counts were standardised to proportions prior to calculation so that dissimilarity reflects relative spatial shifts rather than overall abundance changes.

Lake-specific seasonal maxima were extracted strictly to describe the maximum recorded use of individual lakes. Because maxima across lakes could occur on different dates, their sum was not interpreted as a number of simultaneously active nests or as a population estimate.

### 2.5. Environmental variables

The surface area of each study lake was determined from satellite imagery from July 2025 in Google Earth Pro. The boundary of each lake, including the fringing emergent and floating vegetation used for nesting, was digitised as a polygon by the observer responsible for the field surveys, and surface areas (0.18–2.25 ha) were calculated from the digitised polygons. Lake area measured from the 2025 imagery was treated as a time-invariant proxy for the structural size of each lake, not as a measure of the open-water area available in any particular year.

Lake area was used to test whether variation in lake size was associated with the spatial distribution of Whiskered Tern nests. Other potentially relevant habitat characteristics, including floating vegetation structure, water depth, water-level fluctuations, predation and human disturbance, were not systematically quantified during the study and were therefore not included as explanatory variables.

### 2.6. Statistical analyses

Temporal variation in breeding activity across the study area was examined using the repeated surveys conducted within each breeding season. The total number of active nests recorded per survey period was analysed as count data, with year and sampling date considered as potential explanatory variables.

Seasonal changes in nest abundance were modelled as a function of day of year. Because the relationship between breeding activity and date was not assumed to be linear, generalised additive models (GAMs) were used to model seasonal trajectories: survey-period totals of active nests were related to a smooth function of day of year, with year as a fixed factor (total nests ∼ year + s(day of year, k = 5)). Count distributions were examined for overdispersion, and a negative binomial error distribution was therefore used, with models fitted by restricted maximum likelihood. A model with year-specific seasonal smooths (total nests ∼ year + s(day of year, by year, k = 5)) was compared with the common-smooth model using AIC, with both models refitted by maximum likelihood for the comparison. Residual autocorrelation was inspected visually using within-year autocorrelation functions of the deviance residuals; because survey intervals were not strictly regular, formal fixed-lag autocorrelation tests were not applied.

Differences in the spatial distribution of nests among lakes and years were interpreted using the metrics defined in Section 2.4: lake occupancy, dominant-lake proportion, Simpson’s concentration index (C) and Pielou’s evenness (*J’*), together with Bray–Curtis dissimilarity between years. The association between lake area and breeding activity was evaluated using mixed models with lake identity as a random effect, fitted as GAMs with a random-effect smooth for lake: lake-specific seasonal maximum nest counts were modelled with a negative binomial error distribution as a function of log-transformed lake area and year (seasonal maximum ∼ log(lake area) + year + s(lake, bs = “re”)), and annual lake occupancy (defined as the presence of at least one active nest on a lake in a given year) was modelled separately as a binomial response with log-transformed lake area as predictor (occupancy ∼ log(lake area) + s(lake, bs = “re”)). Year could not be included as a factor in the occupancy model because all nine lakes were occupied in 2022.

Model assumptions were evaluated using appropriate residual diagnostics. Competing model structures were compared using AIC, and effect estimates are reported with standard errors together with sample sizes. Analyses were performed in R 4.4.3 (R Core Team, 2025), using the vegan package (Oksanen et al., 2022) for the diversity, concentration and dissimilarity indices, mgcv (Wood, 2017) for generalised additive (mixed) models, and ggplot2 (Wickham, 2016) for visualisation.

## 3. Results

### 3.1. Interannual redistribution and spatial concentration

Whiskered Tern nest distribution varied substantially among the nine monitored lakes between 2022 and 2025 (Fig. 2). The number of occupied lakes dropped from nine in 2022 (seasonal maximum 32 nests, on Lake 7) to six in 2023 (maximum 25 nests, on Lake 6) and three in 2024 (maximum 29 nests, on Lake 6), before rising again to six in 2025, when breeding activity concentrated heavily on Lakes 4 and 1 (seasonal maxima of 132 and 68 nests, respectively). Per-lake trajectories are shown in Fig. S1.

**Figure 2.**
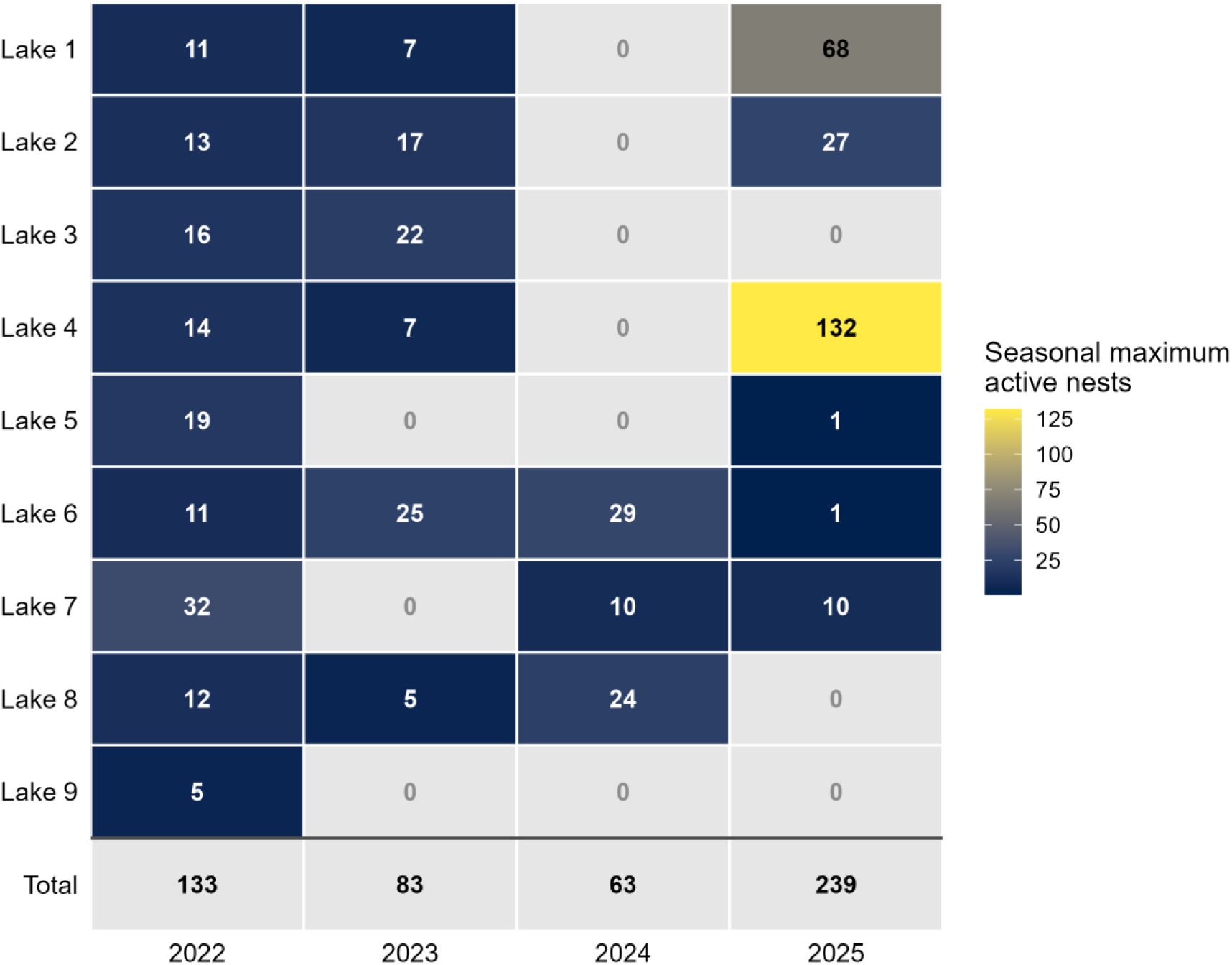
Seasonal maximum number of active Whiskered Tern *Chlidonias hybrida* nests recorded on each of the nine monitored lakes in Văcărești Natural Park, 2022–2025. Cell values are lake-specific seasonal maxima; grey cells denote lakes with no recorded nesting (zero). The bottom row gives the annual sum of the lake-specific seasonal maxima. Because the maxima of different lakes may have occurred on different survey dates, these sums are not numbers of simultaneously active nests and not estimates of breeding population size.

The spatial concentration of breeding activity at the annual peak survey was markedly higher in 2024–2025 than in 2022–2023 (Fig. 3). During 2022–2023, breeding activity was distributed across 5–6 lakes at the annual peak, with the dominant lake holding 41– 46% of nests (C = 0.29–0.32). By 2024–2025, the number of occupied lakes at the peak dropped to 2–4, while the dominant-lake proportion rose to 55–56% (C = 0.42–0.50). This pattern remained unchanged when the alternative peak-survey vector was used for 2022 (Table S4), and a similar increase in concentration was evident when the metrics were calculated from lake-specific seasonal maxima (Tables S2a, S2b). Because these metrics are based on only four annual values, the among-year changes are interpreted descriptively rather than as a formal temporal trend.

**Figure 3.**
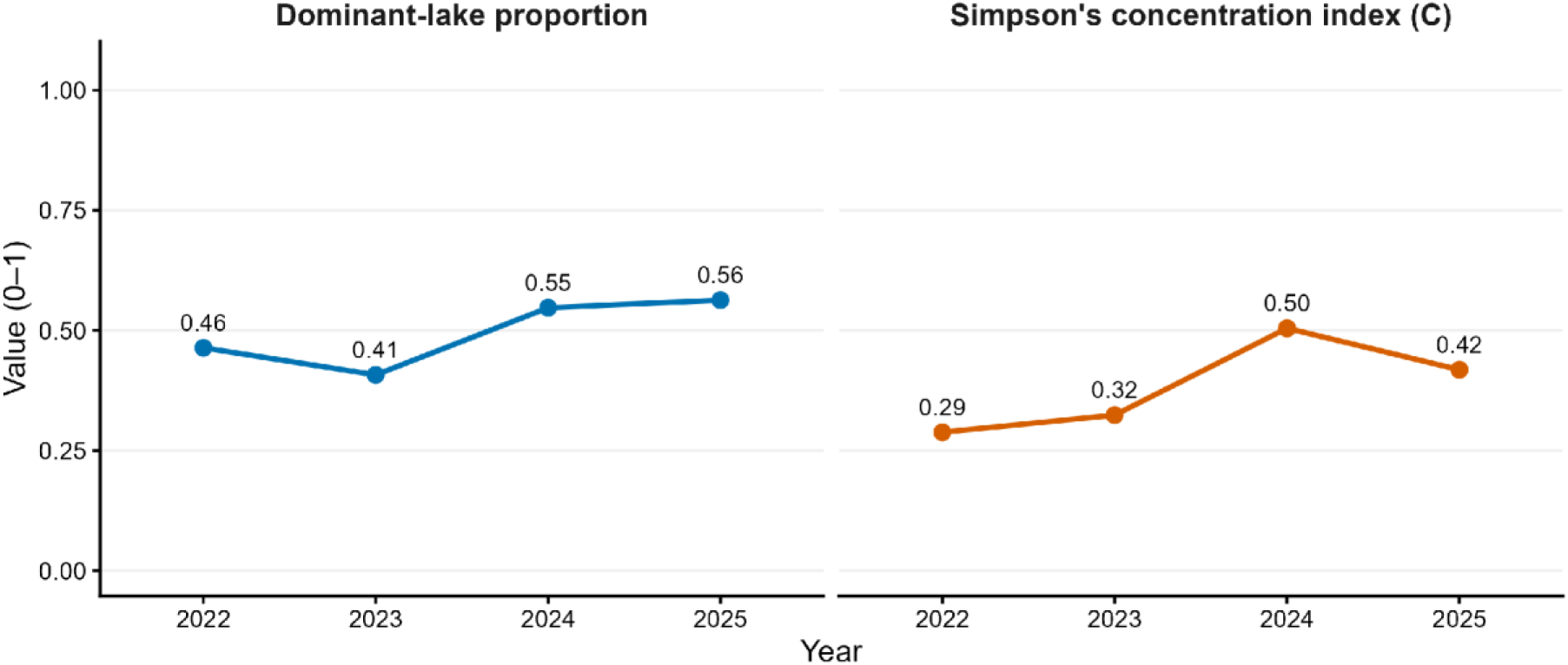
Interannual change in the spatial concentration of Whiskered Tern breeding activity among the nine lakes, 2022–2025, calculated from the simultaneous lake counts of the annual peak survey: the proportion of nests on the dominant lake and Simpson’s concentration index (C = Σ p_i_^2^). Higher values indicate more spatially concentrated breeding activity. As the metrics are based on only four annual values, they are interpreted descriptively rather than as a statistically tested temporal trend.

Pairwise Bray–Curtis dissimilarities based on the simultaneous peak surveys revealed substantial interannual shifts in the spatial distribution of breeding activity (Fig. 4). The spatial distribution was most similar between the consecutive early years (dissimilarity = 0.66 between 2022 and 2023), whereas comparisons involving 2025 showed the highest dissimilarity (≥ 0.86), culminating in complete spatial turnover between the 2024 and 2025 peaks (1.00): the lakes supporting all nests at the 2024 peak (Lakes 6 and 8) held no nests at the 2025 peak, when breeding shifted entirely to Lakes 1, 2, 4 and 7. The same pattern emerged when dissimilarities were calculated from lake-specific seasonal maxima (Table S3); in particular, the primary colonies of 2024 (Lakes 6 and 8; maxima of 29 and 24 nests) were virtually abandoned in 2025, while the newly dominant Lakes 4 and 1 expanded from zero nests to seasonal maxima of 132 and 68 nests, respectively.

**Figure 4.**
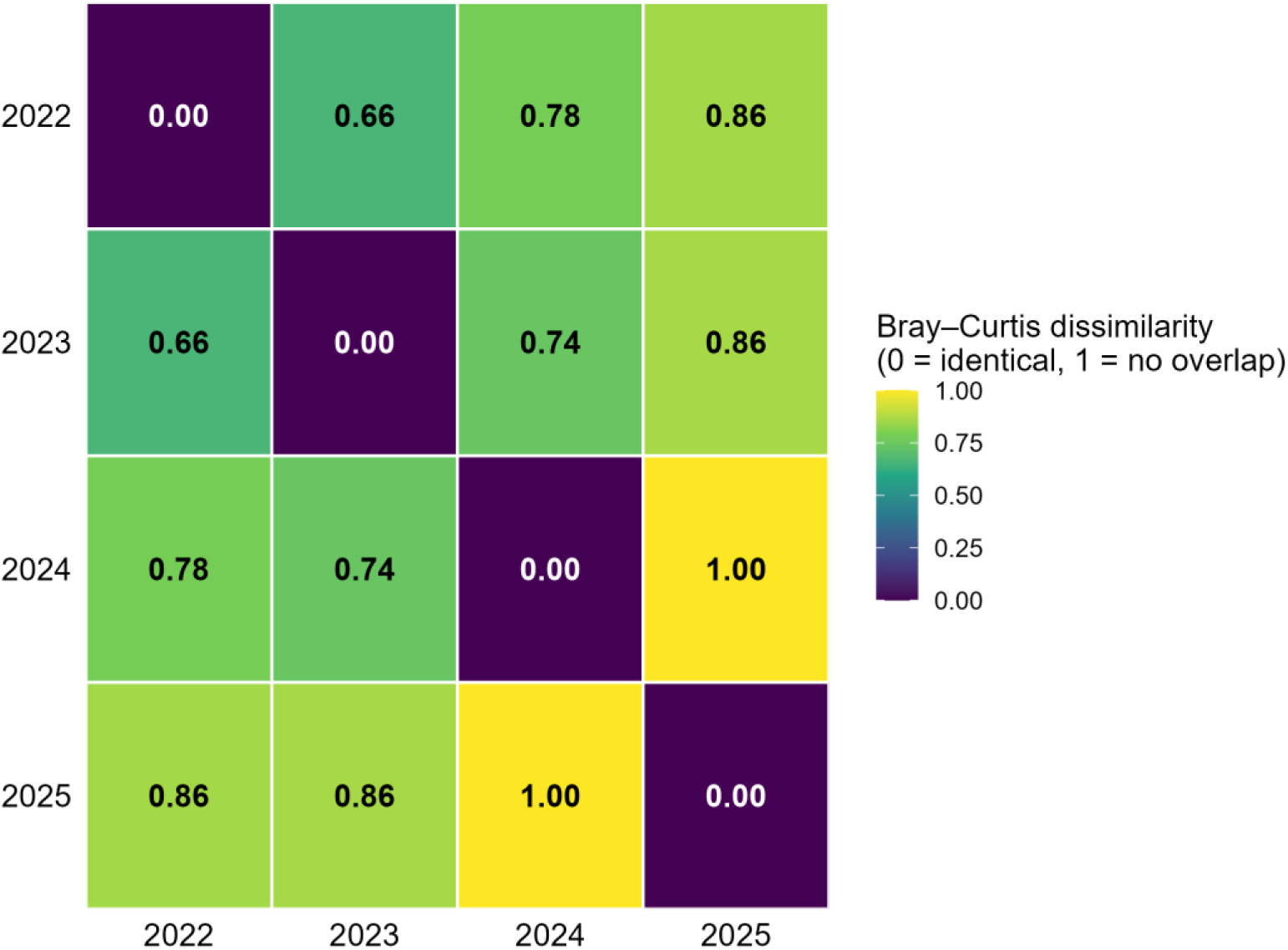
Pairwise Bray–Curtis dissimilarity between the relative lake distributions (proportions of the annual peak total) at the annual peak surveys, 2022–2025. Values range from 0 (identical relative distribution among the nine lakes) to 1 (no overlap in lake use); higher values indicate greater interannual redistribution of breeding activity.

### 3.2. Seasonal dynamics and temporal synchrony

Survey-period totals showed strong seasonal variation in all four years, peaking between late May and mid-June (Fig. 5). Active nests were first detected in mid-May in 2022, 2023 and 2025 (11, 12 and 20 May, respectively), but only in late May in 2024 (28 May). A negative binomial GAM accounted for 82.3% of the deviance (smooth of day of year: edf = 3.8, χ² = 88.3, P < 0.001, n = 51). Overall nest numbers were significantly lower in 2024 than in 2022 (rate ratio = 0.12, P < 0.001), whereas 2023 (rate ratio = 0.53) and 2025 (0.87) did not differ significantly from 2022 (P = 0.19 and P = 0.77, respectively). A model with year-specific seasonal smooths received only weak additional support over the common smooth (ML ΔAIC = 2.7) and yielded much less precise year contrasts; annual rate ratios are therefore reported from the common-smooth model, and year-to-year differences in seasonal trajectories are described qualitatively.

**Figure 5.**
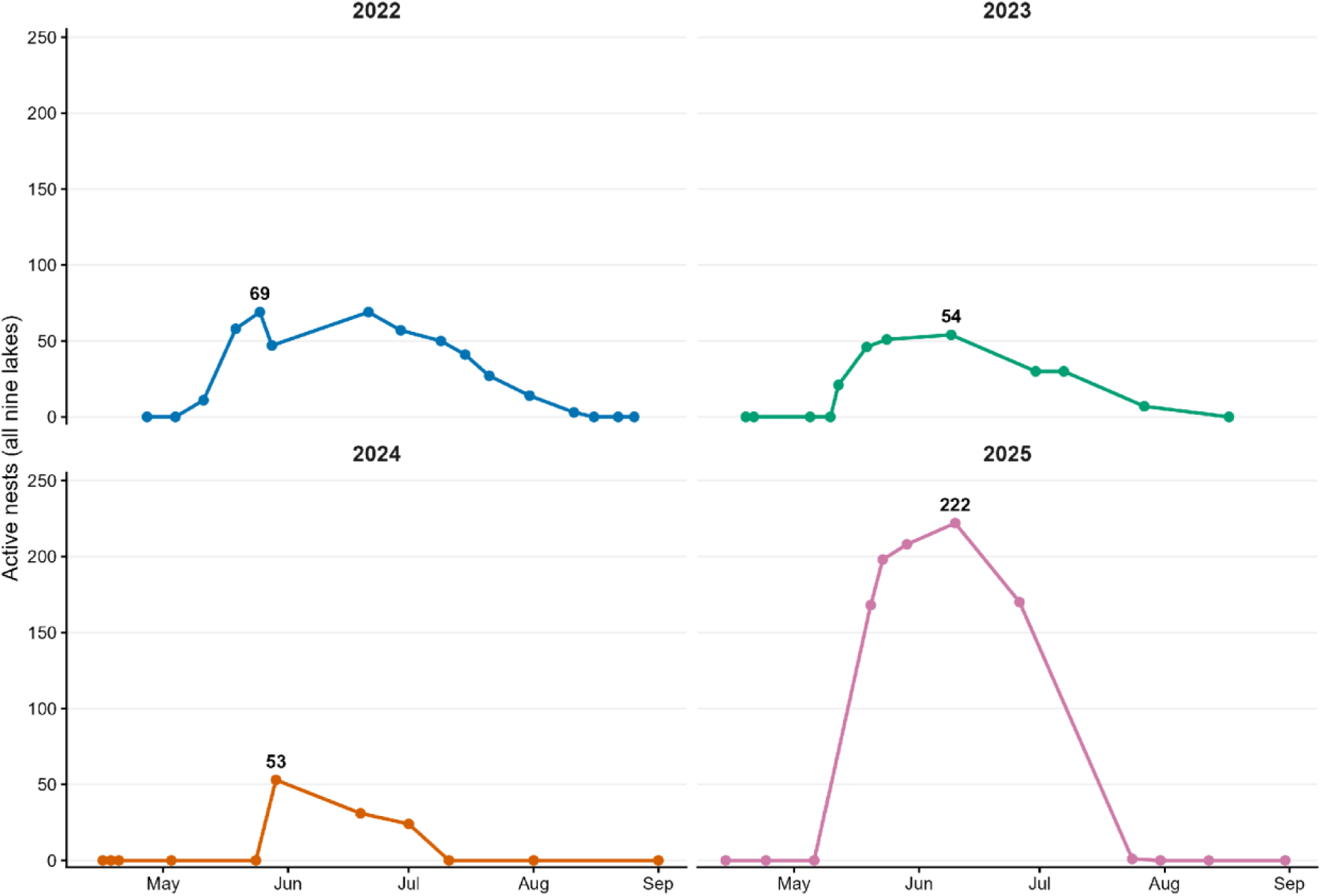
Seasonal dynamics of the total number of simultaneously active Whiskered Tern *Chlidonias hybrida* nests across the nine monitored lakes in each study year. Points show survey-period totals (surveys conducted on consecutive days combined); the annual peak is labelled. All panels share the same y-axis scale.

Annual peaks of simultaneously active nests across the entire complex ranged from 53– 69 nests in 2022–2024 to 222 nests in 2025 (Fig. 5). The degree of temporal synchrony among colonies varied markedly across years. In 2022, the simultaneous peak represented only 52% of the summed lake-specific seasonal maxima (69 vs. 133 nests), reflecting asynchronous colony peaks over an extended nesting period (May to August). In contrast, the simultaneous peaks of 2024 and 2025 reached 84% and 93% of the summed maxima, respectively, indicating high temporal synchrony and a compressed breeding season.

### 3.3. Within-season changes in spatial concentration

Within-season survey trajectories further highlighted the contrast in spatial concentration among years (Fig. 6). In 2022, spatial concentration remained low throughout most of the season (C = 0.17–0.21), with 6–7 lakes holding active nests simultaneously and the identity of the dominant lake shifting repeatedly. A similar within-season turnover occurred in 2023 at intermediate concentration levels (C = 0.25–0.35). In contrast, spatial concentration in 2024 and 2025 remained consistently high throughout the season (C ≥ 0.40 for survey periods with ≥ 10 nests), with a single lake holding the largest colony all season (Lake 6 in 2024, Lake 4 in 2025). Breeding activity thus became both more spatially concentrated and more spatially stable within seasons.

**Figure 6.**
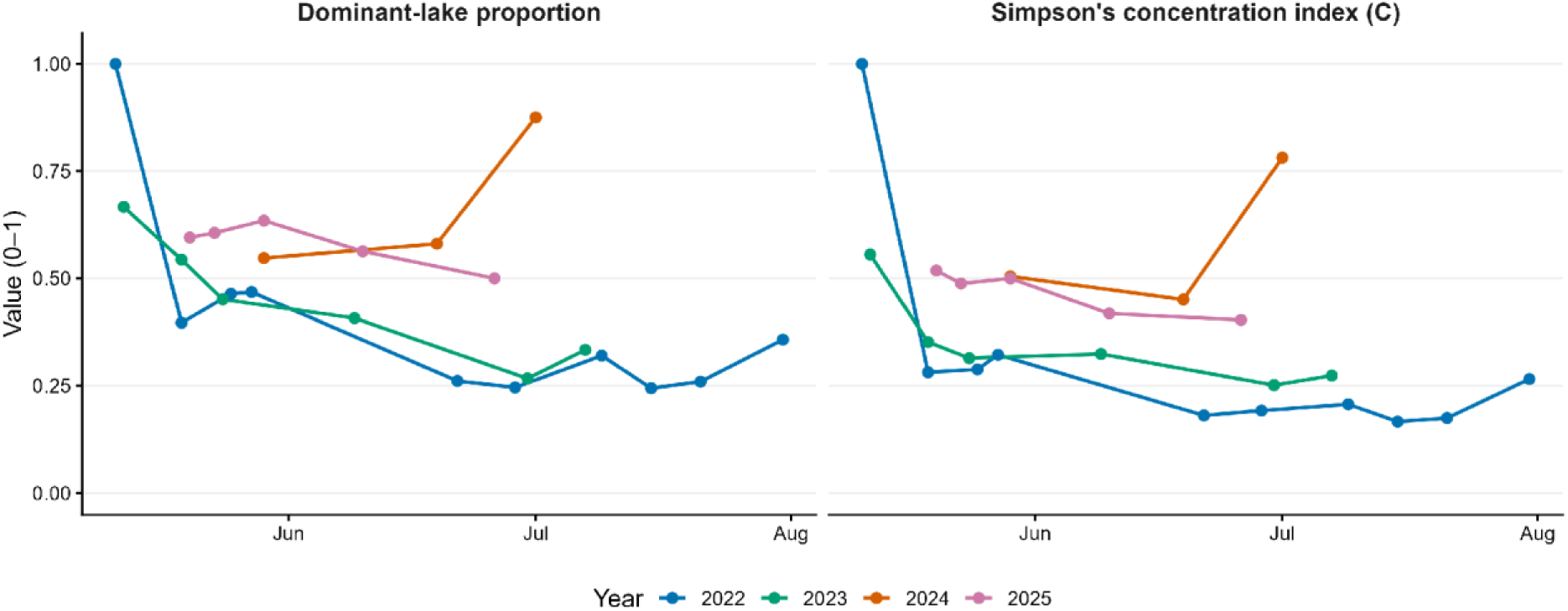
Within-season changes in the spatial concentration of Whiskered Tern breeding activity, shown as the dominant-lake proportion and Simpson’s concentration index (C) for each survey period with at least 10 active nests (a threshold applied to avoid unstable estimates at very small totals), 2022–2025. In 2022, the first survey period with nests (11 May) comprised a single newly established colony, hence values of 1.0.

### 3.4. Lake area and breeding activity

Lake surface area alone (0.18–2.25 ha) was not detectably associated with the spatial distribution of breeding activity across the nine lakes (Fig. S2). Neither lake-specific seasonal maximum nest counts (negative binomial mixed model: log-area, b = 0.06 ± 0.39, P = 0.88) nor annual lake occupancy (binomial mixed model: b = 0.37 ± 0.44, P = 0.41) was related to lake size. Likewise, the rank correlation between lake area and the four-year mean seasonal maximum was weak (Spearman’s ρ = 0.23, n = 9, P = 0.55). Notably, the largest single breeding aggregation occurred on one of the smaller lakes (Lake 4, 0.70 ha, 132 nests in 2025), whereas the largest lake (Lake 1, 2.25 ha) supported substantial numbers only in 2025.

## 4. Discussion

Over just four breeding seasons (2022–2025), Whiskered Tern nest distribution in Văcărești Natural Park reorganised substantially. In 2022, breeding was spread across all nine lakes, with asynchronous, shifting colony peaks throughout an extended season. By 2024–2025, nesting had contracted onto fewer lakes, dominated by a single site with highly synchronised colony peaks. Although four annual values preclude formal trend testing, the consistent directional pattern across several independent concentration metrics indicates a genuine spatial reorganisation rather than an index artefact. The among-year contrast was paralleled within seasons: breeding shifted from an asynchronous, multi-centred and temporally extended pattern in 2022 to temporally overlapping breeding centred on a single, seasonally stable lake in 2024–2025. Lake surface area alone explained neither nest numbers nor occupancy, and the largest breeding aggregation of the study formed on one of the smaller lakes. This is consistent with the ecology of the species, in which colony placement tracks the extent and quality of floating vegetation and the prevailing water regime rather than the size of the water body (Paillisson et al., 2006; Ortiz Lledó et al., 2018; Cieślińska et al., 2025). We could not quantify floating vegetation cover, water depth or water-level fluctuations during the study, so the proximate drivers of the observed redistribution remain to be identified, and, given the small number of lakes and the time-invariant area measure, the lake-area analysis should be regarded as exploratory. The entire lake complex lies well within the foraging range of breeding Whiskered Terns from a single colony (2–3 km; Paillisson et al., 2026), so the nine lakes are best viewed not as separate foraging territories but as a dynamic local network of alternative nesting sites, among which breeding activity redistributes between years (Devlin et al., 2008; Wilkinson and Jodice, 2022). Such pronounced colony relocation aligns with the established expectation that *Chlidonias* terns nesting in dynamic marsh habitats exhibit low colony-site tenacity (McNicholl, 1975; Ławicki et al., 2011; Gula and Downs, 2026). The complete spatial redistribution observed between 2024 and 2025 (Bray–Curtis dissimilarity = 1.0 at the annual peak), marked by the total abandonment of the two primary 2024 sites and the colonisation of two previously unoccupied lakes that then hosted the largest colonies, represents an extreme case of site lability. Although the proximate drivers of these shifts cannot be confirmed without systematic microhabitat monitoring, the development of an extensive carpet of suitable nesting substrate (Great Yellow-cress) on Lake 4 appears to have facilitated the aggregation of 132 active nests at a single site in 2025. High spatiotemporal variability in substrate availability may underpin these marked interannual shifts in site selection across the lake network. Indeed, nesting in our study area occurred on a broad range of substrates, from filamentous green algae to various vascular plants, many of which display pronounced interannual fluctuations in coverage and spatial extent. Crucially, the study area lacks large, stable stands of perennial floating-leaved plants, such as White Water-lily, Yellow Water-lily *Nuphar lutea*, Fringed Water-lily *Nymphoides peltata* or other predictable aquatic vegetation like Water Chestnut *Trapa natans*. Adapted to highly unpredictable breeding conditions in water levels and aquatic vegetation, the Whiskered Tern and its congeners are widely regarded as nomadic (Gochfeld et al., 2020). This nomadic strategy involves widespread prospecting, often across hundreds of kilometres during spring migration (Vágási et al., 2016), to evaluate local breeding conditions; when encountering suitable conditions, individuals rapidly aggregate, occasionally forming exceptionally large breeding concentrations. Because birds in our study were not individually marked, we cannot distinguish between the redistribution of returning breeders and individual turnover via immigration and emigration. Nevertheless, the rapid build-up to 222 simultaneously active nests in 2025 may partly reflect immigration, a scenario plausible given the demographic expansion and high connectivity of eastern European populations (Ledwoń et al., 2014; Dayton et al., 2017).

The studied population also showed considerable plasticity in breeding phenology, ranging from asynchronous, protracted nesting attempts (2022) to highly compressed and synchronised breeding peaks (2024–2025). This variation may reflect spatiotemporal heterogeneity in the timing of suitable microhabitat availability across lakes and years (Paillisson et al., 2006), combined with marked spatial differences in the rate of new nest initiation among individual colonies (Ledwoń M. unpublished data).

Conspecific attraction provides one plausible mechanism for the observed concentration of breeders on a single lake: birds settling where conspecifics are already present would make an occupied site grow at the expense of alternatives (Danchin and Wagner, 1997). Performance-based habitat copying (Danchin et al., 1998) could act in the same direction, although this cannot be evaluated without lake-specific information on reproductive success. The persistence of Lake 6 as the dominant site in 2023–2024, followed by the shift to Lake 4 in 2025, is compatible with such self-reinforcing settlement, although this cannot be tested without individual marking and lake-specific data on reproductive success. Aggregated breeding with closely coinciding colony peaks may bring anti-predator benefits such as predator swamping, although these benefits are conditional on the local predator community (Ims, 1990; Descamps, 2019). Concentration also carries an evident risk: with more than half of all nests on a single small lake, one localised event (a water-level change, a predator or human disturbance) could affect a large proportion of the park’s active breeding attempts at once.

Our results illustrate the conservation value of spontaneously developed urban wetlands (Kowarik, 2011; Alikhani et al., 2021). Just over three decades after its abandonment, and less than a decade after its designation as a natural park (Anastasiu et al., 2017), Văcărești Natural Park supports a Whiskered Tern breeding aggregation of over two hundred simultaneously active nests in the middle of a capital city. However, the high spatial concentration observed in 2024–2025 makes the aggregation more vulnerable, and its dependence on floating vegetation makes the maintenance of stable water levels during the nesting season (May–July) a likely management priority (Ortiz Lledó et al., 2018). Because previously abandoned lakes were repeatedly re-occupied during the study, a pattern typical of colonial waterbirds breeding in dynamic networks of colony sites (Wyman et al., 2014), management and monitoring should cover the whole lake network, including currently unused lakes and the barely accessible Lake 10, rather than only the currently dominant colony site.

The main limitations of this study are its four-year span, the small number of lakes, the absence of individually marked birds, and the lack of systematic data on reproductive success, vegetation and hydrology; the lake-area analysis is exploratory. At the same time, the compact nine-lake system, surveyed in full every one to three weeks, allows colony-site dynamics to be followed at the scale of an entire local network of nesting sites, which is rarely feasible in extensive natural wetlands. Continued standardised monitoring, combined with individual marking and the quantification of vegetation and water-level dynamics, would allow the mechanisms proposed here (tracking of nesting-substrate availability and conspecific attraction) to be tested directly. Aerial (drone-based) checks of the single pond that cannot be inspected from the ground would close the one remaining gap in spatial coverage.

## 5. Conclusions

Whiskered Tern breeding distribution within the compact wetland network of Văcărești Natural Park reorganised rapidly over four breeding seasons, with pronounced changes in the number of occupied lakes, in the identity and dominance of the main colony lake, and in the temporal synchrony of breeding. The complete spatial turnover between the 2024 and 2025 peak-survey distributions shows that lakes unused in one year can host the largest colonies of the next. Because the redistribution of breeding activity was not detectably related to lake size, and nesting substrates varied strongly among years, the value of individual lakes for this species is best regarded as transient. Although vegetation and water levels were not quantified systematically, the pronounced year-to-year turnover in nesting substrates suggests that the availability of suitable floating vegetation, shaped by local hydrological conditions, is a plausible driver of lake choice and should be a priority for future monitoring. Conservation and monitoring of colonial marsh terns in wetland systems that offer several alternative nesting sites, including urban wetlands, should therefore maintain and follow the entire network rather than focus exclusively on the currently largest colony.

## Acknowledgements

We thank the staff of the Văcărești Natural Park Administration for supporting the fieldwork.

## CRediT authorship contribution statement

Andrei Daniel Cotoară: Conceptualization, Investigation, Data curation, Writing – original draft, Writing – review & editing. Mateusz Ledwoń: Conceptualization, Methodology, Validation, Writing – review & editing. Relu Constantin Giucă: Writing – review & editing. Emil Todorov: Investigation, Resources, Writing – review & editing. Alexandru-Cătălin Birău: Resources, Investigation, Writing – review & editing. Gergely Osváth: Conceptualization, Methodology, Formal analysis, Visualization, Writing – original draft, Writing – review & editing.

## Ethics statement

The study was purely observational; surveys were conducted with the support of the Văcărești Natural Park Administration, from distances that minimised disturbance to breeding birds.

## Declaration of generative AI in scientific writing

During the preparation of this work, authors used a generative language tool (Gemini, Google) to improve the language and readability of parts of the text. The authors reviewed and edited the content and take full responsibility for the content of the published article.

## Data availability

The nest-count data and the R scripts used for all analyses are available from the corresponding authors on reasonable request.

