## Supplementary Material (Tables S1-S4, Figures S1-S3) for "Rapid interannual reorganisation of Whiskered Tern *Chlidonias hybrida* breeding colonies within an urban wetland network"

**Table S1.** Survey effort and the annual peak number of simultaneously active Whiskered Tern *Chlidonias hybrida* nests on the nine monitored lakes in Văcărești Natural Park, Bucharest, 2022–2025. Survey periods merge complementary visits on consecutive days; the single incomplete visit in 2025 was excluded (see Methods of the main text).

| **Year** | **First – last survey** | **Survey periods (n)** | **Median interval (days)** | **Peak simultaneous nests (date)** |
|---|---|---|---|---|
| 2022 | 27 Apr – 26 Aug | 16 | 7 | 69 (25 May) |
| 2023 | 19 Apr – 17 Aug | 12 | 7 | 54 (9 June) |
| 2024 | 15 Apr – 31 Aug | 11 | 12 | 53 (28 May) |
| 2025 | 14 Apr – 31 Aug | 12 | 12 | 222 (10 June) |

Table S2a. Annual metrics of the spatial concentration of Whiskered Tern breeding activity, calculated from the simultaneous lake counts of the annual peak survey (primary analysis in the main text).

| Year | Lakes with nests at peak | Dominant proportion | Simpson C | Pielou J′ |
|---|---|---|---|---|
| 2022 | 6 | 0.46 | 0.29 | 0.82 |
| 2023 | 5 | 0.41 | 0.32 | 0.76 |
| 2024 | 2 | 0.55 | 0.50 | 0.99 |
| 2025 | 4 | 0.56 | 0.42 | 0.74 |

Table S2b. The same metrics calculated from the lake-specific seasonal maxima (descriptive comparison). Occupied lakes (annual) is the number of lakes with at least one active nest during the year.

| Year | Occupied lakes (annual) | Dominant proportion | Simpson C | Pielou J′ |
|---|---|---|---|---|
| 2022 | 9 | 0.24 | 0.14 | 0.95 |
| 2023 | 6 | 0.30 | 0.22 | 0.91 |
| 2024 | 3 | 0.46 | 0.38 | 0.93 |
| 2025 | 6 | 0.55 | 0.40 | 0.62 |

Table S3. Pairwise Bray–Curtis dissimilarity between years calculated from the lake-specific seasonal maxima, standardised to proportions of the annual sum of maxima (the corresponding values based on the annual peak-survey vectors are shown in Fig. 4 of the main text).

| | 2022 | 2023 | 2024 | 2025 |
|---|---|---|---|---|
| **2022** | – | 0.47 | 0.67 | 0.66 |

| | | | | |
|---|---|---|---|---|
| **2023** | 0.47 | – | 0.64 | 0.71 |
| **2024** | 0.67 | 0.64 | – | 0.95 |
| **2025** | 0.66 | 0.71 | 0.95 | – |

Table S4. Sensitivity of the annual peak-survey comparisons to the tied 2022 peak. In 2022 the annual maximum of 69 simultaneously active nests was reached in two survey periods (25 May and 21–22 June); the main text uses the earlier period. Using the later period instead leaves the among-year pattern unchanged and makes the 2022 distribution even less concentrated.

| **Metric** | **25 May (primary)** | **21–22 June (alternative)** |
|---|---|---|
| Lakes with nests at peak | 6 | 7 |
| Dominant proportion | 0.46 | 0.26 |
| Simpson C | 0.29 | 0.18 |
| Pielou J′ | 0.82 | 0.94 |
| Bray–Curtis vs. 2023 | 0.66 | 0.61 |
| Bray–Curtis vs. 2024 | 0.78 | 0.75 |
| Bray–Curtis vs. 2025 | 0.86 | 0.82 |

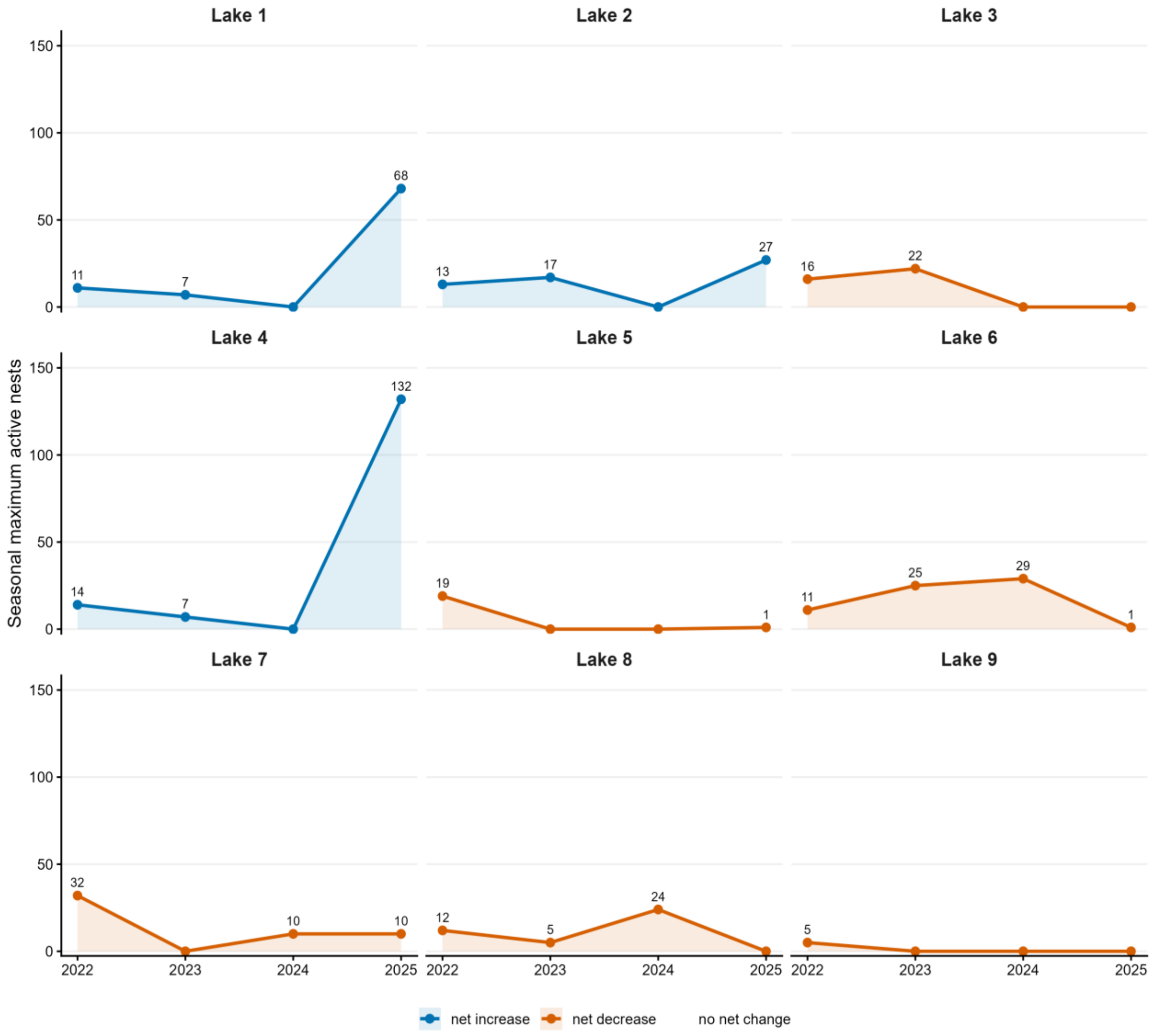

Lake 1
Lake 2
Lake 3
Lake 4
Lake 5
Lake 6
Lake 7
Lake 8
Lake 9
Seasonal maximum active nests
0
50
100
150
2022
2023
2024
2025
11
7
68
13
17
27
16
22
14
7
132
19
1
11
25
29
1
32
10
10
12
5
24
5
net increase
net decrease
no net change

**Figure S1.** Per-lake trajectories of the seasonal maximum number of active Whiskered Tern *Chlidonias hybrida* nests, 2022–2025. Each lake is shown in a separate panel; line colour indicates the net change between 2022 and 2025 (blue = net increase, orange = net decrease, grey = no net change). Note the contrasting patterns, e.g. the marked increases on Lakes 1 and 4 in 2025 and the declines on several previously used lakes (e.g. Lakes 3, 5, 6 and 9).

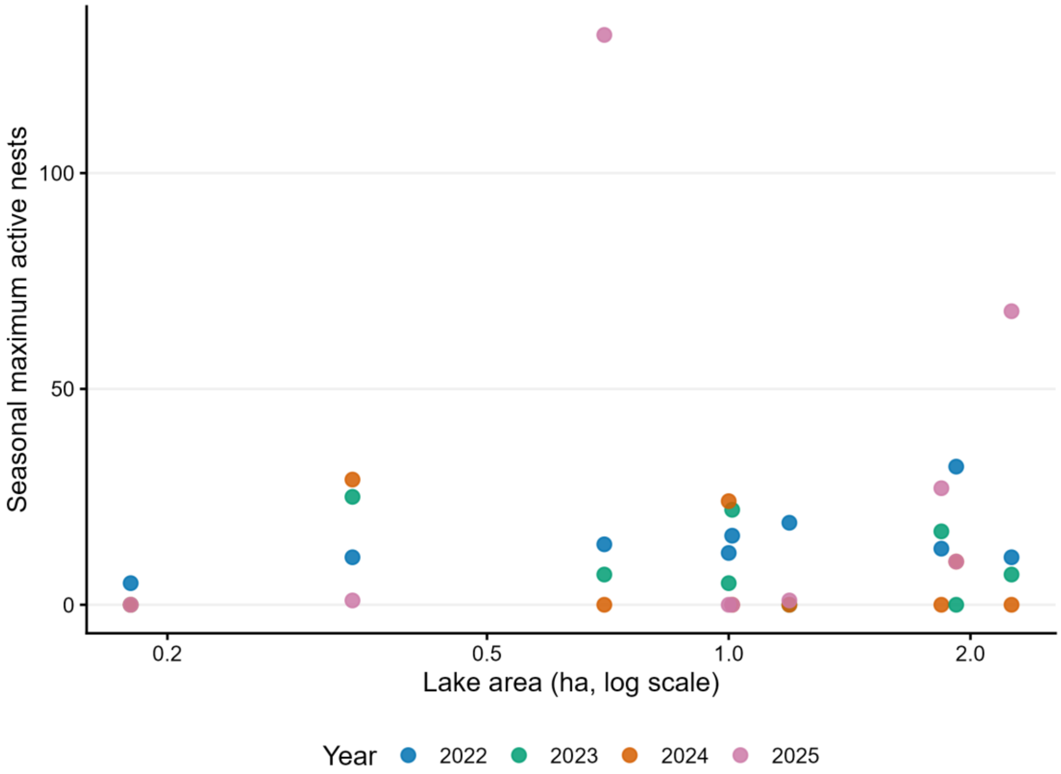

Figure S2.Lake-specific seasonal maximum numbers of active Whiskered Tern nests in relation to lake area (log scale), 2022–2025. Lake areas were measured from July 2025 satellite imagery and treated as a time-invariant structural proxy for lake size; neither nest numbers nor lake occupancy was significantly related to lake area (see main text).

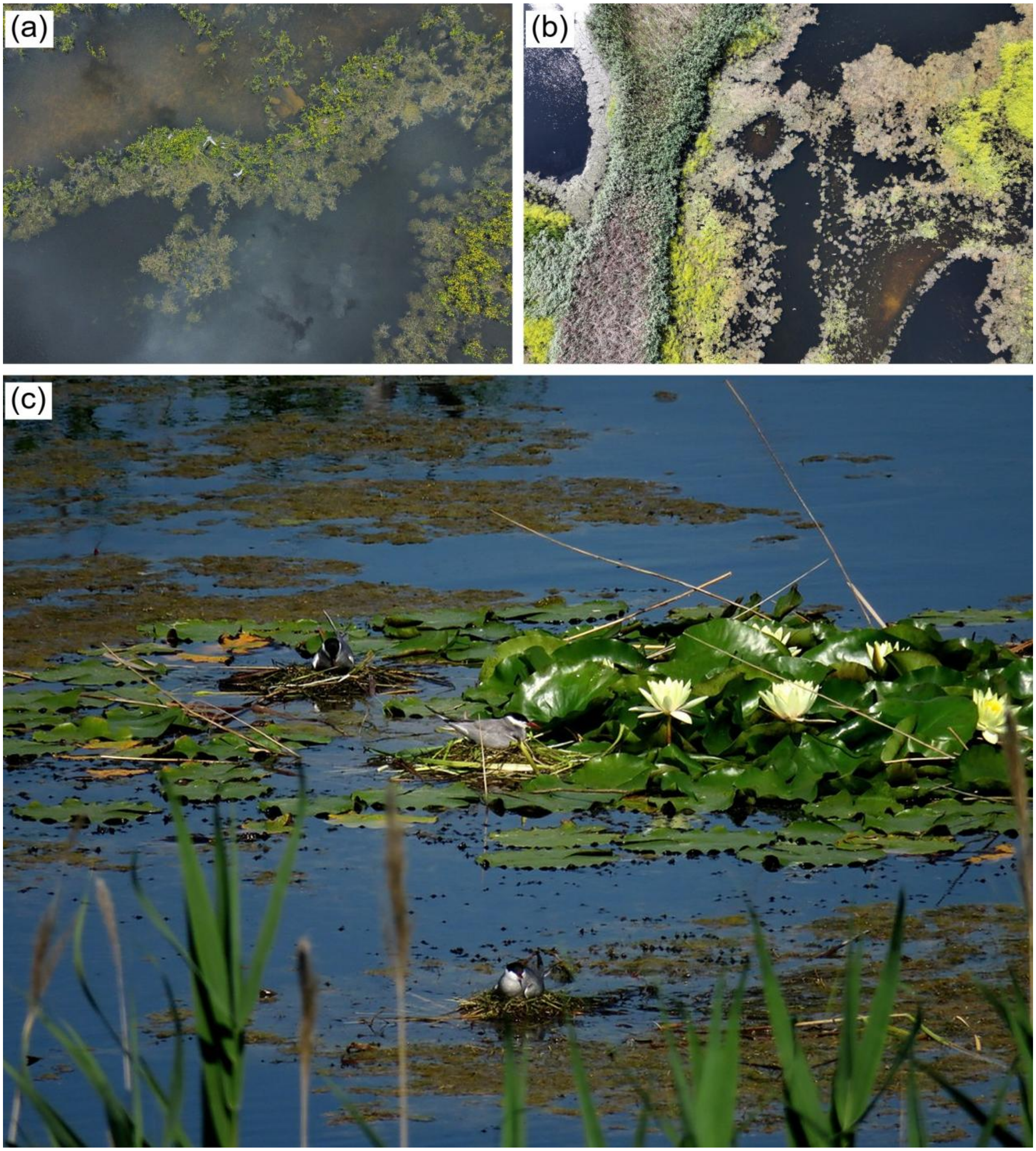

Figure S3. Nesting habitat and breeding Whiskered Terns *Chlidonias hybrida* in Văcărești Natural Park. (a) Part of the Lake 4 colony settling on a flowering carpet of Great Yellow-cress *Rorippa amphibia* (drone image, 6 May 2025); (b) aerial view of the vegetation mosaic of the monitored lakes (drone image, 15 May 2025); (c) incubating adults on floating nests among White Water Lily *Nymphaea alba*. Photographs: (a, b) Emil Todorov; (c) Andrei Daniel Cotoară.
